# PB-kPRED: knowledge-based prediction of protein backbone conformation using a structural alphabet

**DOI:** 10.1101/127423

**Authors:** Iyanar Vetrivel, Swapnil Mahajan, Manoj Tyagi, Lionel Hoffmann, Yves-Henri Sanejouand, Narayanaswamy Srinivasan, Alexandre G. de Brevern, Frédéric Cadet, Bernard Offmann

## Abstract

Libraries of structural prototypes that abstract protein local structures are known as structural alphabets and have proven to be very useful in various aspects of protein structure analyses and predictions. One such library, Protein Blocks (PBs), is composed of 16 standard 5-residues long structural prototypes. This form of analyzing proteins involves drafting its structure as a string of PBs. Thus, predicting the local structure of a protein in terms of protein blocks is a step towards the objective of predicting its 3-D structure. Here a new approach, kPred, is proposed towards this aim that is independent of the evolutionary information available. It involves (i) organizing the structural knowledge in the form of a database of pentapeptide fragments extracted from all protein structures in the PDB and (ii) apply a purely knowledge-based algorithm, not relying on secondary structure predictions or sequence alignment profiles, to scan this database and predict most probable backbone conformations for the protein local structures.

Based on the strategy used for scanning the database, the method was able to achieve efficient mean Q_16_ accuracies between 40.8% and 66.3% for a non-redundant subset of the PDB filtered at 30% sequence identity cut-off. The impact of these scanning strategies on the prediction was evaluated and is discussed. A scoring function that gives a good estimate of the accuracy of prediction was further developed. This score estimates very well the accuracy of the algorithm (R^2^ of 0.82). An online version of the tool is provided freely for non-commercial usage at http://www.bo-protscience.fr/kpred/.

## Introduction

Knowledge of protein structure considerably helps towards understanding protein function. The Protein Data Bank (PDB) that serves as the central repository of knowledge for the protein structural biology community contains more than 125,000 protein structures and its growth has been considerable in the past decade^1^. This number is however still far below the ∼70 million protein sequences referenced in UniProt database^2^. Hence it is at stake to find methods to bridge this considerable gap. Computational methods for predicting protein secondary and tertiary structure have persistently tried to fill it. In this paper, we explore the ability of a structural alphabet based prediction method to fulfill in part this role.

Since the seminal works by Kabsch and Sander in 1984^3^, one of the most popular and rewarding computational method to predict and analyze protein structures is by breaking them down to their constituent parts in the so-called fragment-based approach. Multiple fragment libraries have been developed so far and they differ in the number of fragments, the length of the fragments, the methods used for clustering and the criteria used for clustering. The first fragment library was developed by Unger and co-workers^4^. There are reviews that give a good overview of the different fragment libraries developed since then^5,6^. Also referred to as structural alphabets (SAs), these have shed some light on the sub-secondary structure level intricacies in proteins^7^. By identifying redundant structural fragments found in proteins, structural alphabets help in abstracting protein structures accurately. Such collections of fragments have also been used in methods that attempt to reconstitute protein structures^8-10^.

In that respect, a SA called *protein blocks* (PBs) was developed for the purpose of describing and predicting the local backbone structure of proteins^11,12^. This SA accounts for all local backbone conformations in protein structures available in the Protein Data Bank (PDB). Since then, PBs have been used in various applications^11^: for structural motif identification^13–15^, structural alignments^16,17^ and fold recognition^18,19^. There have also been various efforts to use PBs to predict protein local structure. These approaches are based on the Bayes theorem^9,12^ support vector machines^20–22^ and neural networks^23^. Some of these methods have used prior predictions of classical three state secondary structures (svmPRAT^22^ uses YASSPP^24^, SVM-PB-Pred^21^ uses GOR^25^ and Etchebest et. al.^26^ use PSI-PRED^27^) and sequence alignment profiles like position specific scoring matrices (PSSMs) are used by LOCUSTRA^20^, SVM-PB-Pred^21^ and Dong and coworkers methodology^23^. The currently available web-based tools that can predict local structure in terms of protein blocks are LocPred^28^ and SVM-PB-Pred^21^. The former implements a Bayesian methodology and the latter is SVM-based.

In this work we describe PB-kPRED, a purely fragment and knowledge-based approach to predict local backbone structure of proteins in terms of protein blocks and a web-based tool that implements the method. In essence, it takes no other inputs than the amino acid sequence of a query and interrogates a database of pentapeptides extracted from protein structures, without using evolutionary information. It returns the predicted local structures of the polypeptide chain in the form of a sequence of protein blocks. Very importantly, PB-kPRED also implements a scoring function that efficiently auto-evaluates the quality of the prediction.

## Methods

### Dataset

All the protein chains from PDB^1^ were segregated into clusters culled at 30% sequence identity using the BLASTClust algorithm^28^ resulting in a collection of 15,544 clusters. The dataset which we set-up comprises of 15,554 protein chains each corresponding to the best representative structure available from each of these clusters and is hereafter termed as “PDB30 dataset”. Preference was given to crystallographic structures over NMR and electron microscopy structures and also preferring better resolution and lowest R-value structures. Out of these 15,544 structures, 14,207 are crystallographic structures, 1,128 are from NMR experiments and 209 are solved by electron microscopy. Further, chains smaller than 100 residues were filtered out. We preferred to keep the NMR and EM structures, as we wanted to investigate if the experimental method impact on the quality of the predictions. For each of these 15,544 proteins the subsets of PDB that were homologous at 30%, 40%, 50%, 70%, 90%, 95% and 100% as reported by BLASTClust were also calculated in order to implement the “*Hybrid method with noise filtering*” scheme.

### Protein Blocks

The set of protein blocks (PBs) is a structural alphabet composed of 16 structural prototypes each representing backbone conformation of a fragment of 5 contiguous residues^11,12^. The 16 PBs are represented by the letters a to *p* and were identified from a collection of 228 non-redundant proteins. Clustering these pentapeptides was based on the 8 dihedral angles (ψ_i−2_, φ_i−1_, ψ_i−1_, φ_i_, ψ_i_, φ_i+1_, ψ_+1_, φ_i+2_) that define their local backbone conformation. An unsupervised learning algorithm (Kohonen algorithm) was used to arrive to an unbiased classification of the dihedral vectors and to the definition of standard dihedral angles for each PB. Protein blocks are assigned on the basis of the dissimilarity measure called root mean square deviation on angular values (*rmsda*) between observed dihedral angles and the standard dihedral angles for the 16 PBs. The PB with lowest *rmsda* is assigned to the central residue of the pentapeptide region. The choice of fragment size as 5 and library size as 16 for the PBs was because 5 consecutive residues capture well the local contacts in regular secondary structures (α-helices and β-strands) and 16-library size is a good balance between the specificity and sensitivity of predictions^12^.

All the 15,544 protein chains from PDB30 dataset were encoded into their corresponding protein blocks sequences (PB sequences) after comparing their backbone φ and ψ torsion angles with the corresponding standard torsion angles for the 16 PBs^11^ using an in-house developed Perl script. Sequence of PBs as observed in crystal and NMR structures were later used as a reference to assess the accuracy of predicted PB sequences.

### Database of pentapeptide conformations from protein structures

A database of pentapeptide conformations (PENTAdb) was developed using known 3-D structures of proteins. PENTAdb is essentially the entire structural information contained in the PDB, broken down into chunks of pentapeptides. A sliding window of 5 residues was used to extract structural features for every overlapping pentapeptide of a polypeptide chain. The dihedral vector associated with the five consecutive residues that is required to assign PBs as described in the previous section was obtained from the DSSP^29^ program. All the information was stored as a MySQL relational database. PENTAdb is maintained up-to-date; the update frequency corresponds to the weekly updates of PDB. The protein chains from which the pentapeptides are extracted are filterable at 30%, 40%, 50%, 70%, 90%, 95% and 100% sequence identity thresholds.

### Prediction scheme

The overall scheme for predicting the local structure in terms of PBs is based on querying the PENTAdb database for every constitutive pentapeptides of a query protein sequence using a sliding window of 5 residues (Figure 1a). Hits from the database are reported as predicted protein blocks (PBs). Predicted PBs are assigned to the central residue of each query pentapeptide. The prediction results are presented at different levels of refinement. The prediction in the coarsest form consists of the list of all the possible PBs for a particular pentapeptide of the query protein sequence. This is the case when multiple hits from PENTAdb database are obtained for a particular query pentapeptide (Figure 1b). The multiple hits correspond to the different conformations, which the pentapeptide has been seen to adopt in protein structures (Figure 1b). When the query pentapeptide is not found in PENTAdb, the information available for the tetrapeptides covering the first four residues with a wildcard for the fifth position was used (Figure 1b) to identify the list of possible PBs with first 4 amino acid residues matching this query. The position of wildcard did not influence the outcome of the results (data not shown). The list of hits thus obtained is referred as *all possible PBs*. This list serves as a framework from which the most probable PB sequence is predicted.

**Figure 1.**
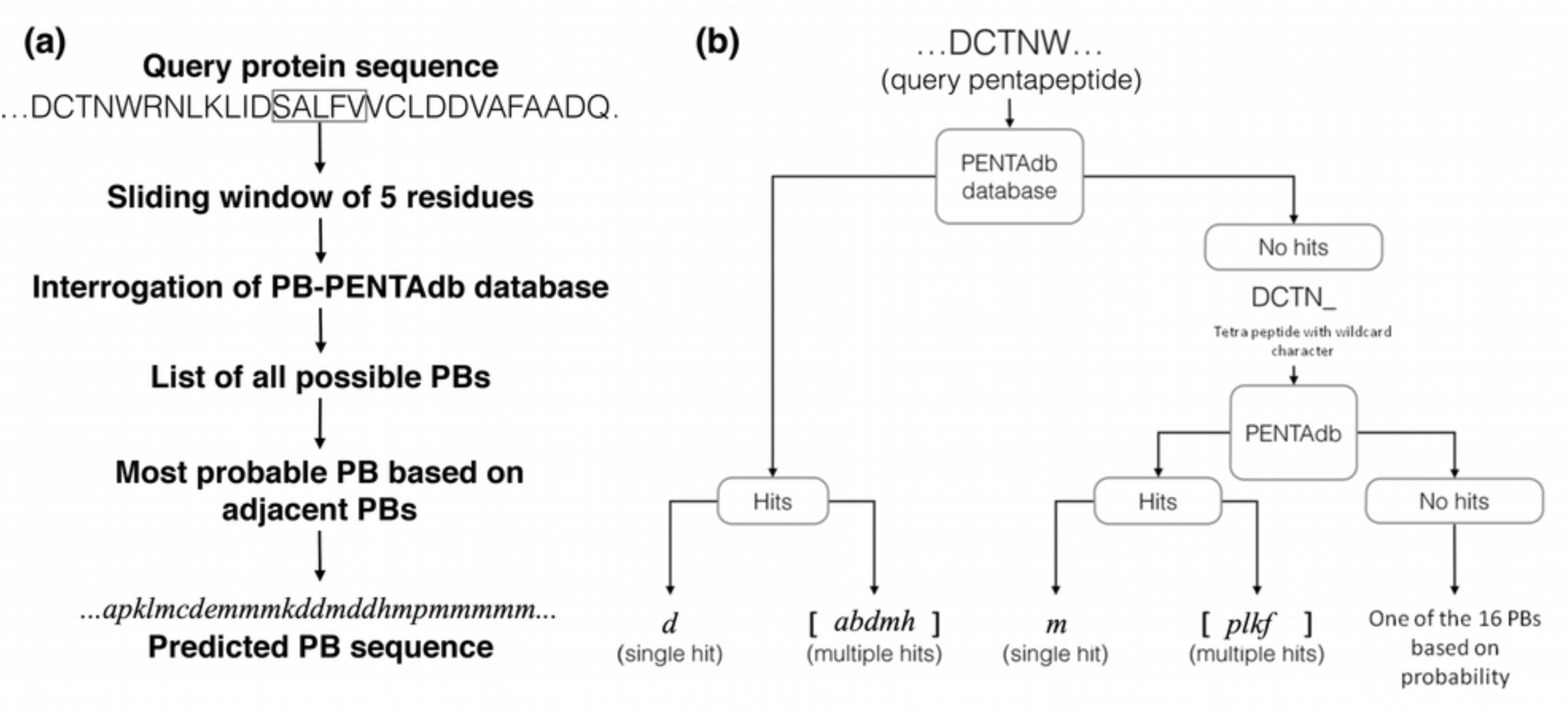
The knowledge-based methodology behind PB-kPRED. (a) Overview of the scheme followed by PB-kPRED for the prediction process. (b) The different outcomes possible when PENTAdb database is queried for a pentapeptide sequence: hits are reported as a single PB or multiple PBs.

Two methods were explored to predict the optimal PB sequence within the list of all the possible PBs obtained after querying the database. The first method, termed as *majority rule method*, is purely probabilistic and consists of simply picking up the most frequently observed PB for each query pentapeptide. As shown in Figure 2, it corresponds to the PB that has highest S1 score, where S1 scores are simply the raw counts of all possible PBs reported by PENTAdb database for the query pentapeptide. In cases when there is no decisive majority (two or more equi-probable PB), both of them are reported as predictions.

**Figure 2.**
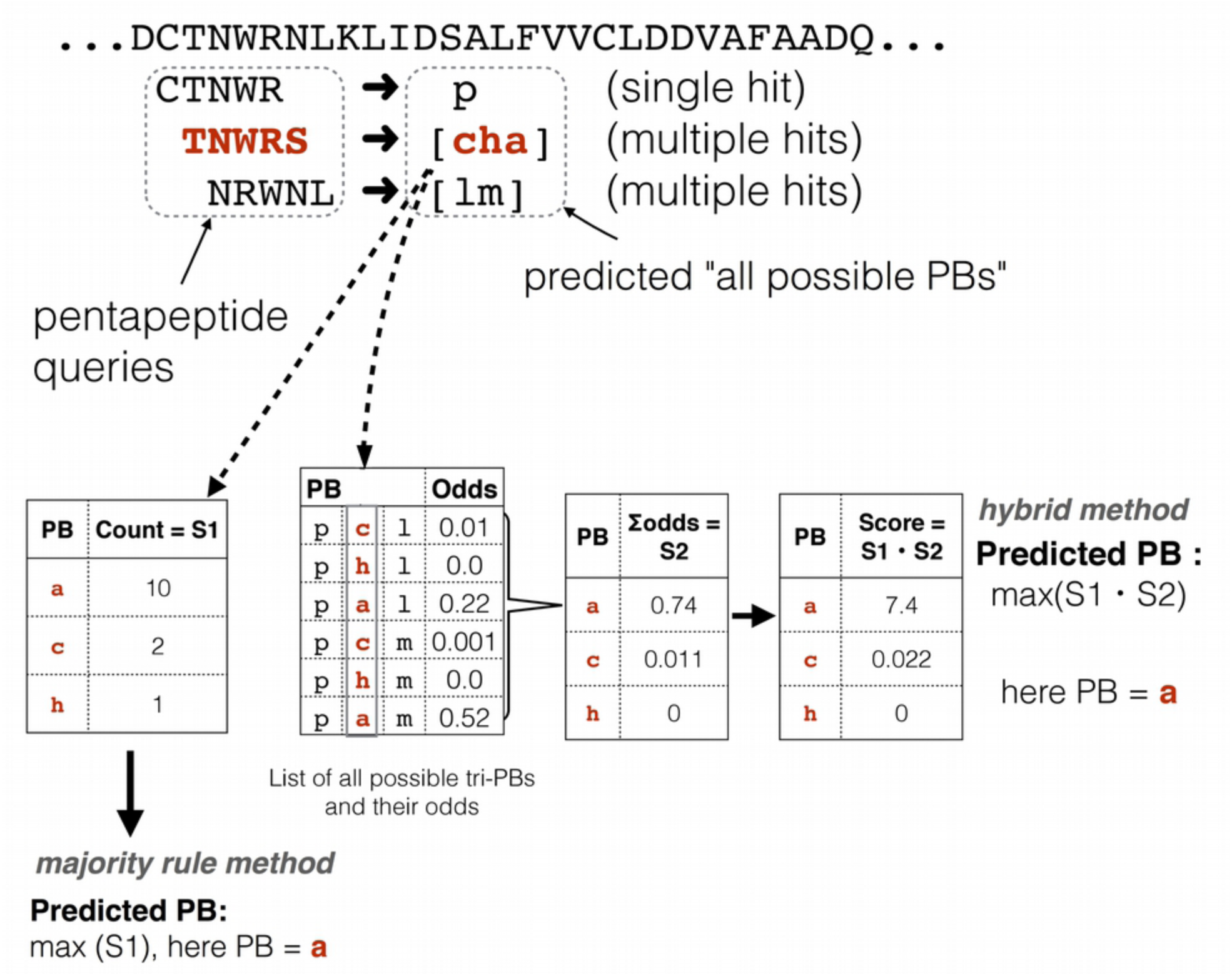
Details of the scoring schemes underlying the *majority rule method* and the *hybrid method*. S1 scores are simply the raw counts of all possible PBs reported by PENTAdb database for a given query pentapeptide. S2 scores are calculated through the summation of the odds of tri-PBs that have a common PB in the central position. For the majority rule method, predictions are based only on the ranking of the S1 scores. For the *hybrid method*, predictions are based on the ranking of the product of scores S1 and S2.

However, it is known that the structure adopted by a short peptide can be highly dependent on its local environment^3^. A second method that integrates contextual information was hence developed and is hereafter termed as *hybrid method*. Here, to predict the local structure of a pentapeptide, the information about the structural status (in terms of PBs) of the two immediately adjacent and overlapping pentapeptides (preceding and succeeding) is also taken into account (see Figure 2). It requires a normalized frequency look-up table for observed motifs of 3 consecutive PBs also termed as tri-PBs (see “additional methods” section of the supplementary material). For each query pentapeptide, in complement to the calculated S1 score, an additional S2 score is calculated as follows. A list of all possible combinations of three successive PBs (tri-PB motifs) is built. This is derived from the list of *all possible PBs* for the query pentapeptide and for its two adjacent pentapeptides (Figure 2). For each possible tri-PB motif, their normalized frequencies (“odds” in Figure 2) are looked up in the tri-PB normalized frequency table. S2 scores are calculated through the summation of the odds of tri-PB motifs that have a common PB in the central position (Figure 2). The predicted PB for the query pentapeptide is determined after multiplying S1 scores by their corresponding S2 scores and taking the highest value among these products (Figure 2). This approach is called the *hybrid method* because it combines the *majority rule method* with contextual information in the prediction process.

### Evaluating PB-kPRED using different subsets of PENTAdb

Two evaluation schemes were developed to benchmark the PB-kPRED methodology. As mentioned above, the query dataset used here constituted of the 15,544 proteins from the PDB30 dataset. The schemes relied on the ability to control which subsection of PENTAdb will be accessible to the prediction algorithm for every query. For example, allowing only pentapeptides in PENTAdb from non-homologues to be accessible by the prediction algorithm emulates a scenario of attempting to predict the local structure of a protein with no homologue of known structure used. On the other hand, as in the case of other local structure prediction methods^12,20–22,26^, it can be advantageous to have the ability to privilege information from homologous structures when these are available to predict the local structure of a query protein. Such a scheme can be emulated by allowing only pentapeptides in PENTAdb from closest detectable homologues to be accessible by the search algorithm.

In first instance, the prediction methodology was assessed with increasing sequence identity cutoffs ranging from 30%, 40%, 50%, 70%, 90%, 95% to 100%, named experiments A1-A8 (see Figure 3a). This scheme is subsequently termed as “*without noise filtering scheme*”.

**Figure 3.**
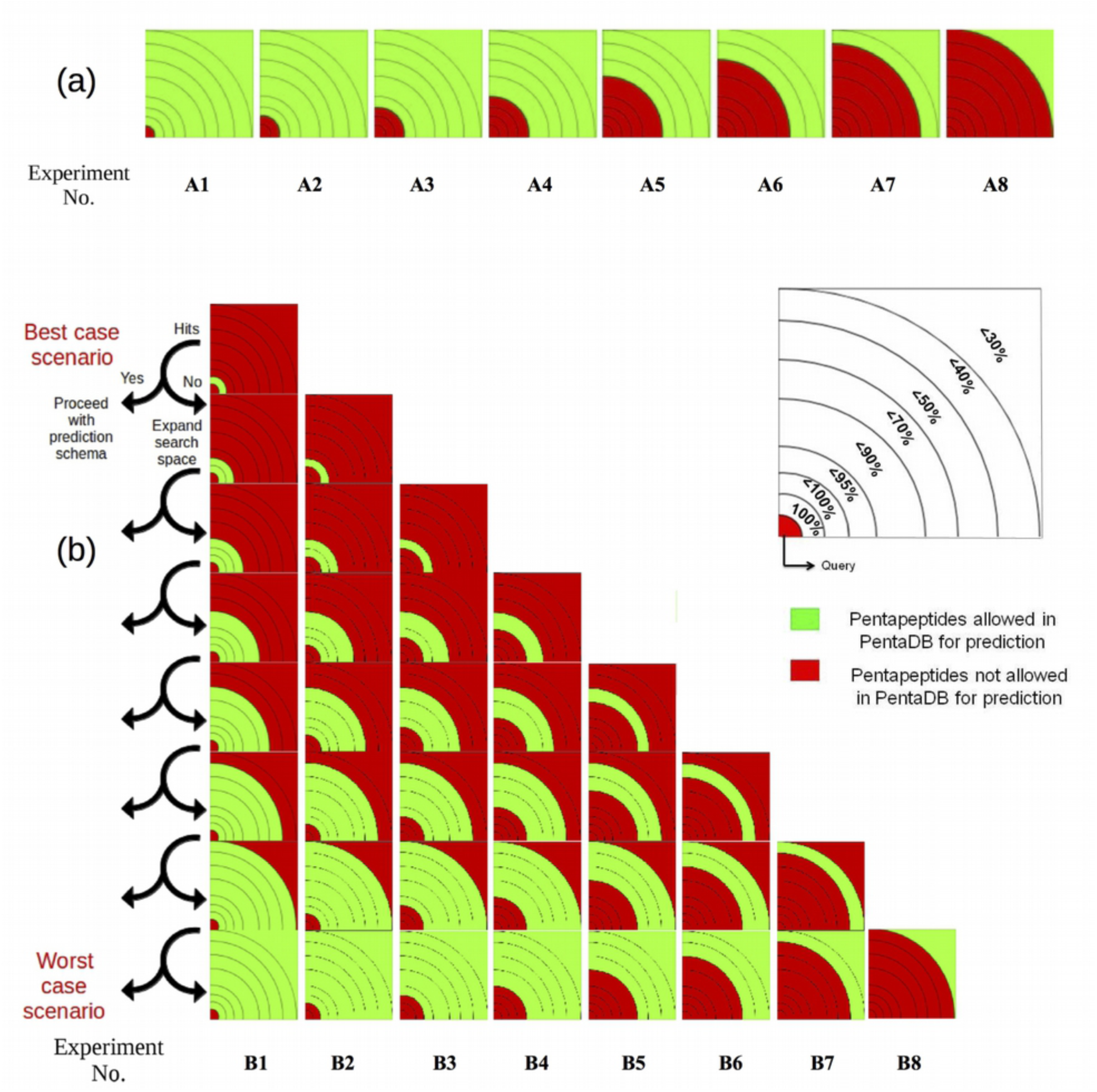
Diagrammatic representations of the schemes used by PB-kPRED for querying PENTAdb with (a) representing the so-called “classic” or “without *noise filtering scheme*” and (b) representing the “with *noise filtering scheme*”. Sections of the database accessible are indicated in green and those not accessible in red. The sections are delimited by sequence identity thresholds. In both schemes, eight different experiments (A1 to A8 and B1 to B8) were performed. See “additional methods file” in supplementary material for detailed legend of this figure.

In second instance, an alternative assessment scheme hereby called the “*with noise filtering scheme*” was applied to further assess the PB-kPRED methodology (experiments B1-B8, see Figure 3b). It aimed at evaluating how privileging information from close homologues, when available, contributed to the quality of the predictions. In brief, the algorithm initially searches for a pentapeptide among the closest homologues first. If the search finds a hit, then the hit is used for the prediction; otherwise the search space is increased to include the immediately next level of more distantly related homologues. This process is repeated until a hit is obtained. Due to this process of introducing more distant homologues in a conditional fashion, wrong pentapeptides (noise) from PENTAdb were potentially filtered out, hence the name *with noise filtering scheme*. In all the cases, care was taken to exclude the pentapeptides from the query proteins themselves.

Reducing the PB predictions into a binary outcome permits the use of classical Mathews correlation coefficient (MCC) to compare our predictions to a random choice. MCCs for the 16 PBs were evaluated based on a confusion matrix similar. For each PB, MCC was calculated according to Equation 1.

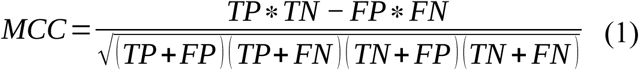

### A scoring function to estimate the accuracy of the predictions

A probabilistic scoring function was developed for the *a posteriori* analysis of the predicted PB sequence through namely the analysis of its content in penta-PB motifs, with the objective of providing a measure of how accurate PB-kPRED was performing. The principle of the analysis relies on the fact that not all penta-PBs are commissioned by proteins at the same frequency. Indeed, many successions of 5 consecutive PBs are highly improbable because they are geometrically not allowed as explicated by the Ramachandran rules. The probabilistic function is hence based on the look-up table of normalized frequencies of successive penta-PB motifs observed in a non-redundant set of protein structures (see “additional methods” in supplementary material). In brief, using a sliding window of 5 consecutive PBs (penta-PB motif) along the predicted PB sequence, the normalized frequencies of all penta-PB motifs were looked-up in the penta-PB frequency table. The logarithm of these normalized frequencies were then summed and divided by the length of the predicted PB sequence to generate an *accuracy score* (*A*) as shown here:

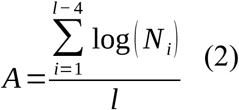
where *A* is the *accuracy score* for a predicted PB sequence, *l* is the length of the PB sequence, *N* is the normalized frequency of the penta-PB motif observed at window position *i* in the PB sequence. Since an overlapping sliding window of five consecutive PBs is used, the total number of penta-PB motifs (*i.e* the number of windows) is *l*-4. In the case a particular penta-PB motif has a null value in the frequency table (*i.e* it is never observed), a penalty of −5 was instead added to the score.

## Results

### PENTAdb, a database of pentapeptides from protein structures

A total of 68.84 million pentapeptides obtained from the 0.26 million protein chains and their corresponding local structure represented as one of the 16 PBs were obtained and stored in PENTAdb. Of these 68.84 million pentapeptides, 2.26 million are unique which represents 70.9% of the total number theoretically possible 3.2 million (20^5^) pentapeptides. The content of the database accessible to PB-kPRED at these threshold values is given in Table 1. There is a 32-fold decrease (from 68.62 to 5.13 million) in the number of pentapeptides in PENTAdb when PDB chains not sharing more than 30% sequence identity are considered. Nevertheless, there is only a 1.3 fold decrease in the number of unique pentapeptides present in PENTAdb at this threshold.

**Table 1.** Content of the different subsets of PDB in terms of pentapeptides accessible to PB-kPRED. Shown here are the total number of pentapeptides and unique pentapeptides for the full PDB and for subsets of PDB filtered at different sequence identity cut-off values.

| **Seq. identity cut-off values (%)** | **Total number of chains** | **Total number of unique pentapeptides** | **Total number of pentapeptides** |
| --- | --- | --- | --- |
| <30 | 24,564 | 1,742,890 | 5,126,423 |
| <40 | 28,590 | 1,881,813 | 6,153,846 |
| <50 | 32,588 | 1,985,203 | 7,074,571 |
| <70 | 37,741 | 2,095,663 | 8,336,277 |
| <90 | 42,594 | 2,148,064 | 9,351,888 |
| <95 | 44,714 | 2,157,239 | 9,768,748 |
| <100 | 64,129 | 2,189,924 | 14,791,285 |
| **Full PDB** | **274,920** | **2,268,307** | **68,621,454** |

### Not all possible tri-PB combinations are observed in known protein structures

Out of all the theoretically possible 4,096 (16^3^) tri-PBs, a total of 1,375 (i.e 33.5%) were never observed in the non-redundant PDB30 dataset. Likewise, out of all the 1.04 million (16^5^) theoretically possible penta-PB motifs, only 40,130 (3.8%) were observed in the PDB30 dataset. These results are indicative of the possibility that many combinations of three or five consecutive PBs are stereochemically unfavorable. The distributions of penta-PB motifs at other sequence identity cut-offs i.e. 40%, 50%, 70%, 90%, 95%, 100% and the entire PDB were also computed (Supplementary Table 1). Towards higher sequence identity cut-offs, there was a steady increase in the penta-PB coverage. But this comes at the price of the addition of redundant data. Still the entire PDB covered less than 10% of the total penta-PB space. For calculating the *accuracy score*, the penta-PB frequency table derived from the PDB30 dataset was used even if it contained only 40,130 penta-PB motifs. This might seem to be a small fraction but this was sufficient to efficiently score PB sequences (see below).

### Completeness of PENTAdb for knowledge-based prediction

A quantitative assessment of how often the correct PB can be found in the list of all possible PBs reported for every query protein was performed. This represents the theoretical highest prediction rate attainable for a query protein using the proposed knowledge-based approach. To this end, for every query protein sequence, different portions of the pentapeptide database were made accessible to the prediction algorithm. This is manageable, thanks to the hierarchical clustering at different sequence identity levels by the BLASTClust algorithm^29^. For each of the 15,544 unrelated query protein sequences (PDB30 dataset), only pentapeptides coming from a subset of the PDB that shared sequence identities below an indicated cut-off values (from 30% to 100%) and excluding the query itself were made accessible to PB-kPRED for prediction (see Table 1 for size of the database for each subset). The results are detailed in Table 2. At 30% sequence identity cut-off, the correct PB was found in 71.4% of the case and the success rate increased to 77.3% when only “homologues” sharing 100% sequence identity to the queries were filtered out. When full PDB was used (but excluding the query) as a database, the percentage times the correct PB is in the list of all possible PBs topped to 99.93%. The PB-wise breakdown of these values are further detailed in supplementary Table 2.

**Table 2.** Assessment of the richness of PENTAdb towards knowledge-based prediction of protein backbone in terms of protein blocks. Shown is the percentage of pentapeptide queries for which the correct local conformation was found in the list of all possible PBs reported by the PB-kPRED algorithm after querying PENTAdb. A total number of 15,544 query proteins not sharing more that 30% sequence identity (PDB30 dataset) was used in this assessment.

| Sequence identity cut-off values (%) | Correct PB found in the list of all possible PBs (%) |
| --- | --- |
| <30 | 71.4% |
| <40 | 71.5% |
| <50 | 71.6% |
| <70 | 71.9% |
| <90 | 72.5% |
| <95 | 72.8% |
| <100 | 77.3% |
| FULL PDB* | 99.93% |

\* all PDB chains (FULL PDB) were accessible for prediction by PB-kPRED but excluding the query protein.

### Prediction accuracies

The average prediction accuracies for the PDB30 query proteins using the *majority rule method* and the *hybrid method* using the classic scheme for querying the database are given in Table 3. When homologues sharing ≥30% sequence identity with each of the queries were removed from the database, PB-kPRED performed with an average Q16 accuracy of 39.2% and 40.8% for the *majority rule method* and *hybrid method* respectively (Table 3). Surprisingly, the effect of enlarging the database to include closer homologues sharing ≤95% sequence identity with the queries improved only marginally the prediction accuracies reaching on average 40.4% and 42.4% for *majority rule method* and *hybrid method* respectively. Accuracy topped to 58.0% and 54.6% respectively when full PDB (excluding the query itself) was used as database for prediction. This overall gain in accuracy is due to an incremental increase of accuracy across all the 16 PBs.

**Table 3.** Evaluation of performance of PB-kPRED knowledge-based approach to predict local conformations of protein backbone in terms of protein blocks. Shown are the accuracies for the PDB30 dataset using both the *majority rule method* and *hybrid method*. For each of the 15,544 query protein sequences, the portion of PENTAdb accessible for prediction was dynamically determined using MySQL queries: only pentapeptides coming from protein chains in PDB that shared sequence identities below the indicated cut-off values were accessible to PB-kPRED for prediction of local structures in terms of PBs.

| Experiment number | Sequence identity cut-off values (%) | Majority rule method | Hybrid method without noise filtering |
| --- | --- | --- | --- |
| A1 | FULL PDB* | 58.0%±12.9 | 54.6%±22.0 |
| A2 | <100 | 44.0%±14.7 | 48.0%±18.7 |
| A3 | <95 | 40.4%±12.7 | 42.4%±16.1 |
| A4 | <90 | 40.1%±12.5 | 42.0%±15.9 |
| A5 | <70 | 39.7%±12.5 | 41.4%±15.8 |
| A6 | <50 | 39.4%±12.5 | 41.0%±15.9 |
| A7 | <40 | 39.3%±12.5 | 40.9%±15.9 |
| A8 | <30 | 39.2%±12.5 | 40.8%±15.9 |

\* All PDB chains (FULL PDB) were accessible for prediction by PB-kPRED but excluding the query protein.

As an attempt to improve the prediction rates, the hybrid method was tested using the *noise filtering scheme* for querying the database whereby, for each query pentapeptide, data in PENTAdb only coming from closest homologues was queried first (see Figure 3). Results are detailed in Table 4. When compared to the *without noise filtering scheme* (Table 3), the prediction rates improved to reach a maximum of 66.3%. Interestingly, for experiments B2 to B7 where closest homologues to be queried first are in the range of <40% to <95% sequence identities, the predictions remained high at a level of about 61.6%. Only in experiment B8 the prediction accuracy rate dropped to 40.8%. This experiment is in fact identical to the one featured for <30% threshold shown in Table 3 using the hybrid method and the *without noise filtering scheme* for querying the database.

**Table 4.**
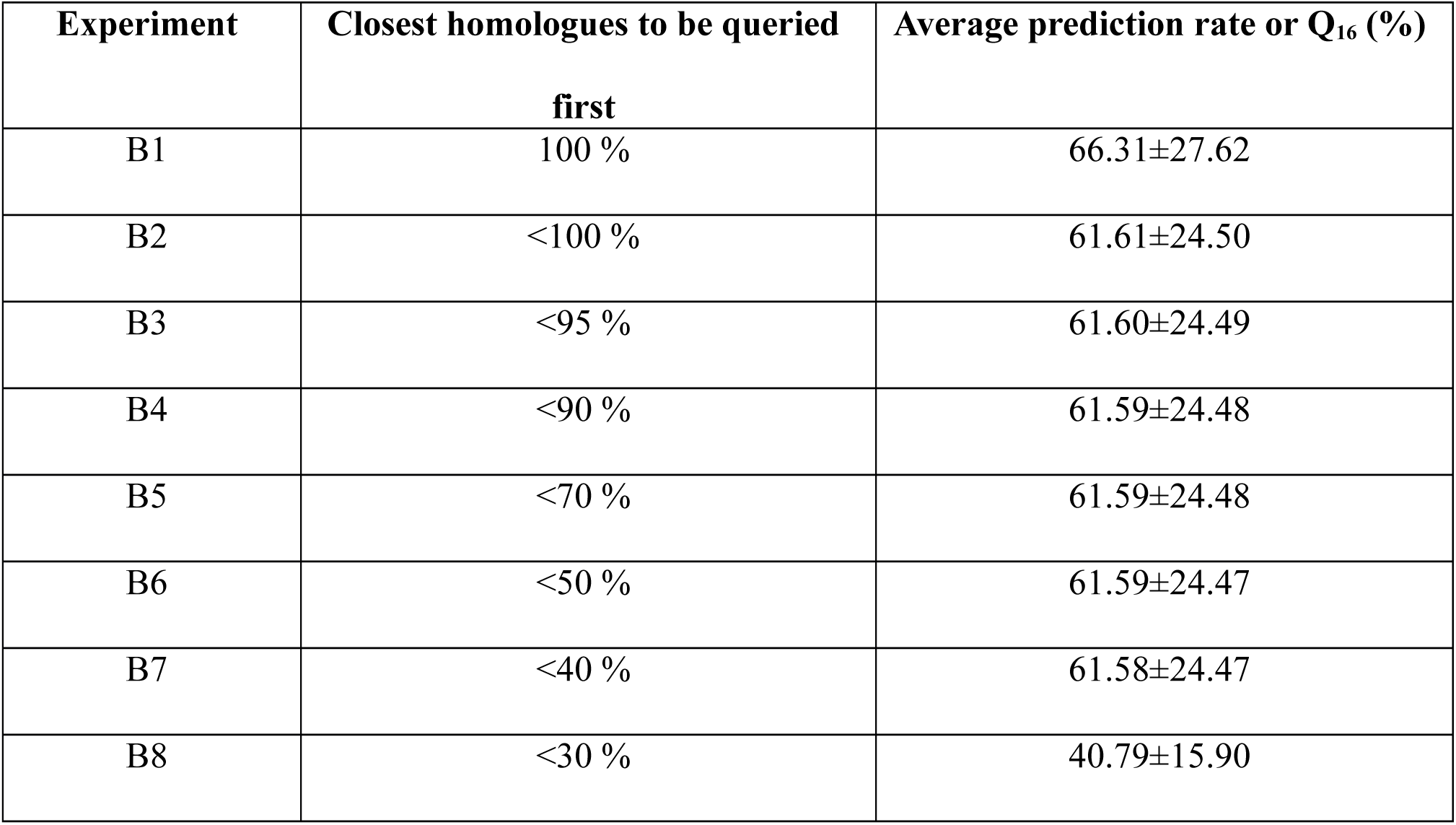
Assessment of the performance of PB-kPRED using the hybrid method with *noise filtering scheme* for querying the database. For each query protein, the portion of the database accessible to the algorithm is first restricted to the closest homologues and if no hits were found, only then the more distant homologues are made accessible progressively. Eight results shown here correspond to the eight experiments represented schematically in Figure 3. Shown are the prediction rates (or Q16) averaged over 15,544 query proteins from the PDB30 dataset that was used in this assessment.

All results further detailed hereafter are concerned with data obtained in experiment B1 where hybrid method was applied using the *noise filtering scheme* for querying PENTAdb and where best predictions were obtained.

The distribution of the prediction accuracies for experiment B1 (see Table 4) shows a bimodal distribution (see Figure 4). A spike in frequency is observed at the >80% range representing the set of queries which have closely related proteins of known structure available in the PDB and for which the method is able to perform extremely well. At the other end of the spectrum, there is an almost normal distribution with an average around the 35%-40% accuracy range. Hence, the mean falls in between these two at 66.31% accuracy. This distribution did not substantially vary when homologues sharing less than 100% to 40% sequence identity to the query corresponding to experiments B2 to B7 respectively (see Figure 3) were queried first (data not shown). However, once the twilight zone of 30% sequence identity is crossed, the accuracy distribution drastically changes to that of a unimodal distribution with a very sharp peak at the 40% range and gradually tapering tail towards the higher accuracies (supplementary Figure 1).

**Figure 4.**
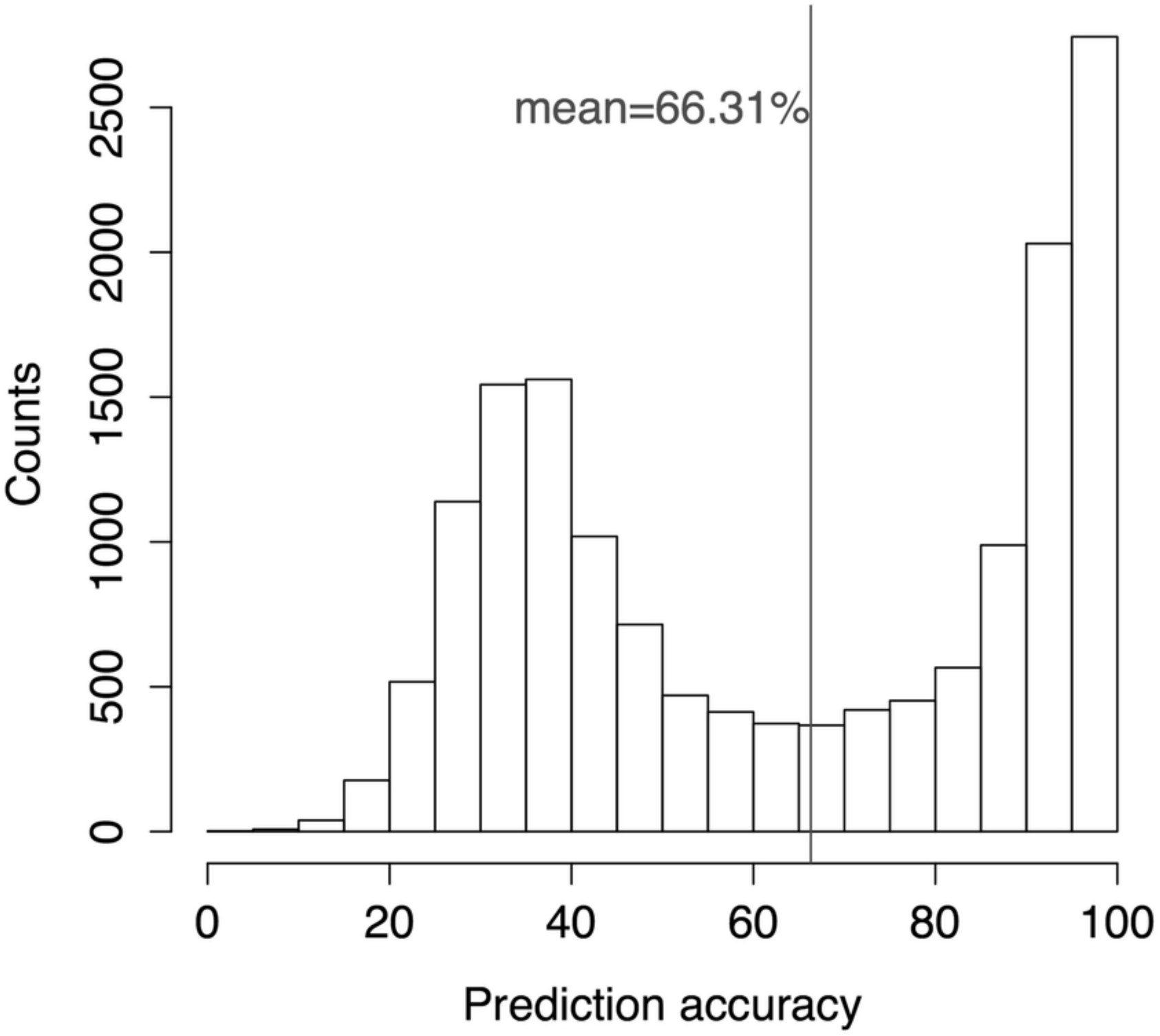
Histogram depicting the distribution of the observed prediction accuracies for 15,544 query proteins by *hybrid method using noise filtering scheme*.

The accuracy by the *hybrid method* using the *noise filtering scheme* was compared to the *majority rule method* (Figure 5). As shown by the data points below the diagonal, the *hybrid method* performed significantly better than the *majority rule method* for a total of 8,195 cases (52.7%) out of the 15,544 protein queries. For remaining 7,245 cases, the *majority rule method* performed slightly better than the *hybrid method*.

**Figure 5.**
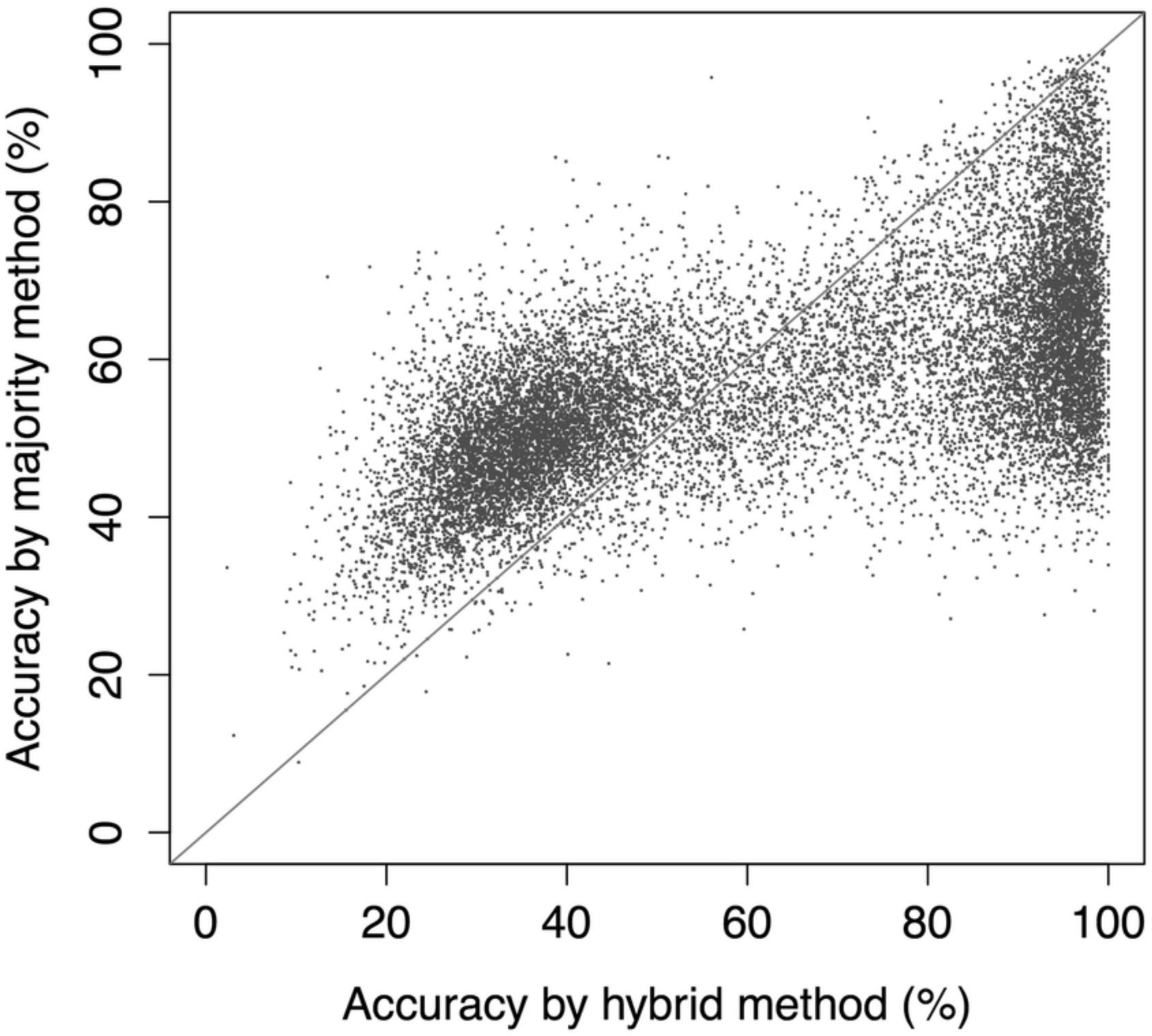
Comparison of the majority rule method without *noise filtering scheme* and the *Hybrid method* with *noise filtering scheme*. Shown are the predictions accuracies for the 15,544 query proteins from PDB30 dataset. The diagonal line separates the points where the majority method performs better and the points where the *hybrid method* performs better. Points lying along the diagonal (bisector) represents the situation where both the methods perform equally.

### PB predictions

Results from the best performing condition (experiment B1 featured in Table 4 and Figure 3b) were further analyzed for the PB-wise prediction rates and compared with published rates from other methods (Table 5). The rates are heterogeneous across the 16 PBs. Top two best-predicted PBs by PB-kPRED were PB *m* and PB *a*, with accuracies of 75.9% and 67.2%, respectively. On the other hand, the two most badly predicted PBs by PB-kPRED were PB *j* and PB *g* with prediction rates of 49.9% and 43.5% respectively. Analysis of the corresponding confusion matrix (see supplementary Table 3) shows that, the prediction algorithm frequently gets confused between the PBs *c* and *d*. PB *c* is wrongly predicted as PB *d* almost 31,000 times (22.4%). The vice-versa, PB *d* being predicted as PB *c* is more than 42,000 times (16.9%). These PBs are in fact highly related (i) as seen from a pure structural point of view (low *rmsda* and similar transitions)^27^ and (ii) as they have been seen to be highly interchangeable thanks to PB substitution matrix^31,32^. As some PBs are highly similar, it is possible to relax the assessment, *i.e*. considering two PB series as equivalent. With such relaxed criteria, the accuracy increases from 66.31% to 68.87% (a 2.56% gain on average). Interestingly, significant increases in accuracies were observed for PB *g* (from 43.5% to 67.4%) and for PB *j* (from 49.98% to 67.2%).

**Table 5.** Assessment of the performance of PB-kPRED and comparison with other previously reported methods. Shown are the PB-wise prediction accuracies for experiment A8 of the majority method and four different experiments of hybrid method with noise filtering. These are compared with PB-wise results from LOCUSTRA^19^ and the method using Bayesian approach developed by Etchebest et al^25^. Experiment B1 PB-wise accuracies were compared with the other two methods and corresponding cell values in bold represent the best accuracy achieved between the experiment B1 of *hybrid method*, LOCUSTRA and Bayes method.

| PBs | PB frequency | Majority method | Hybrid method |  |  |  |  |  | Other methods |  |
| --- | --- | --- | --- | --- | --- | --- | --- | --- | --- | --- |
|  |  | Expt A8 | Expt B8 | Expt B4 | Expt B2 | Expt B1 |  |  | LOCUSTRA | Bayes method |
|  |  | Accuracies |  |  |  |  | Specificity | MCC | Accuracies |  |
| *a* | 3.68% | 34.40% | 45.93% | 64.60% | 64.60% | **67.20%** | 98.15% | 0.69 | 58.16% | 56.60% |
| *b* | 4.29% | 15.19% | 18.99% | 44.52% | 44.58% | **52.15%** | 97.72% | 0.56 | 26.14% | 20.90% |
| *c* | 8.31% | 23.45% | 28.76% | 51.62% | 51.66% | **58.53%** | 95.95% | 0.58 | 44.81% | 32.90% |
| *d* | 18.68% | 39.24% | 40.09% | 61.83% | 61.85% | 67.00% | 94.12% | 0.63 | **71.58%** | 54.00% |
| *e* | 2.18% | 21.78% | 28.59% | 52.30% | 52.38% | **57.45%** | 98.96% | 0.62 | 44.74% | 38.60% |
| *f* | 6.45% | 24.87% | 29.49% | 54.64% | 54.64% | **60.30%** | 97.21% | 0.61 | 41.45% | 30.90% |
| *g* | 1.14% | 10.33% | 14.94% | 36.79% | 36.87% | **43.45%** | 99.19% | 0.51 | 26.84% | 30.10% |
| *h* | 2.10% | 24.25% | 33.02% | 56.90% | 56.89% | **61.05%** | 98.82% | 0.64 | 38.45% | 40.90% |
| *i* | 1.49% | 20.69% | 30.45% | 55.36% | 55.35% | **59.17%** | 99.18% | 0.63 | 36.87% | 38.10% |
| *j* | 0.83% | 15.48% | 20.81% | 42.57% | 42.68% | **49.98%** | 99.40% | 0.56 | 48.19% | 49.70% |
| *k* | 5.25% | 30.46% | 35.59% | 59.30% | 59.32% | **63.93%** | 97.67% | 0.65 | 46.46% | 33.40% |
| *l* | 5.20% | 26.56% | 31.88% | 54.87% | 54.93% | **59.99%** | 97.69% | 0.62 | 42.71% | 35.50% |
| *m* | 32.89% | 60.96% | 55.68% | 72.38% | 72.40% | 75.89% | 91.03% | 0.67 | **83.76%** | 70.60% |
| *n* | 1.78% | 26.48% | 35.40% | 58.16% | 58.21% | **62.15%** | 99.05% | 0.65 | 52.08% | 50.00% |
| *o* | 2.44% | 29.62% | 38.40% | 59.49% | 59.52% | **63.19%** | 98.70% | 0.66 | 55.10% | 48.10% |
| *p* | 3.29% | 23.22% | 31.72% | 54.27% | 54.28% | **59.24%** | 98.25% | 0.62 | 40.80% | 29.20% |

PB-kPRED globally outperformed two other PB prediction methods (see Table 5). Its predictions were better for all the 16 PBs when compared to the Bayes method and better than almost all PBs when compared to LOCUSTRA. Only PBs *d* and *m* were better predicted by this latter method^20^.

A MCC close to +1 indicates a good agreement between the observed and the predicted outcomes and a MCC of close to −1 otherwise. For our analysis all the 16 PBs had MCCs between 0.5 and 0.7. PBs *a* and *m* were close to 0.7, PB *g* at 0.51 and the remaining fluctuated around the 0.6 mark. The sensitivity and specificity ranges were 0.4-0.7 and 0.9-1.0 respectively. A common pattern is observed in the case of PBs corresponding to the regular secondary structure elements (PBs *d* and m): in both these cases, the sensitivity values peak while the specificity values plummet. Although the sensitivity values varied between 0.4 and 0.8, the specificity values were consistently above 0.9 indicating that the method was able to achieve a very high true negative rate.

### Measure of accuracy

A probabilistic scoring function was developed for the *a posteriori* analysis of the predicted PB sequences so as to provide a measure of how accurate the *hybrid method* using the *noise filtering scheme* was performing. An assessment of the scoring function is provided in Figure 6. It shows that the score is correlated with the accuracy of the prediction with a Pearson’s correlation coefficient of 0.82 (Figure 6a). The two distinct clusters of data points correspond to those featured in the histogram in Figure 4. As a further assessment of the scoring function, the scores for the predicted PB sequences were compared with the scores for the actual PB sequences (Figure 6b). It shows that in case of more accurate predictions (rates above 60%), the two scores correlated very well (red points along the diagonal in Figure 6b) with both score values mostly ranging between +1 and +3. In the case of less accurate predictions (rates below 60%), the two scores were no more correlated (green dots below the diagonal in Figure 6b) and scores for predicted PB sequences ranged mostly between −2 and +1.

**Figure 6.**
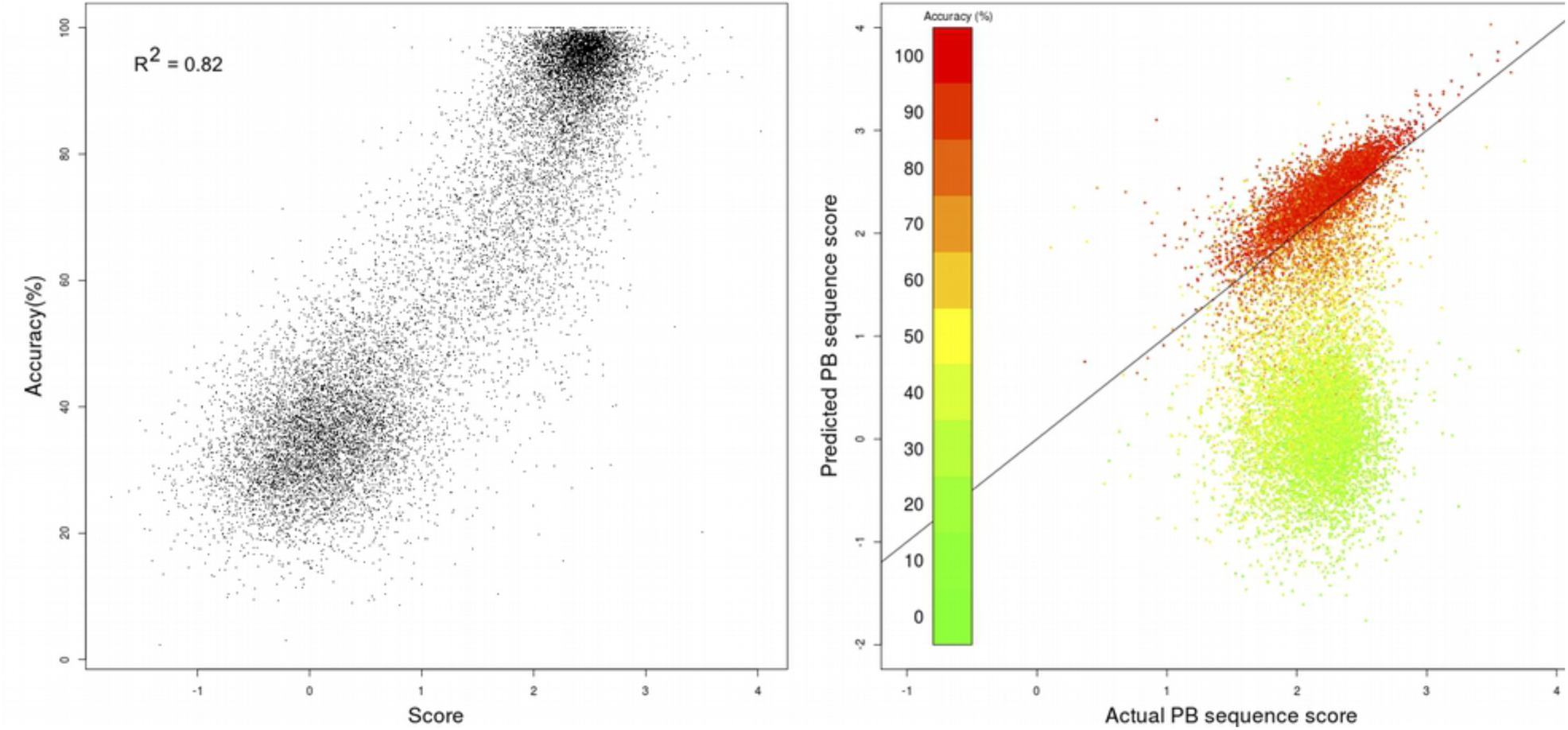
Assessment of the ability of the scoring function to estimate the prediction accuracy of PB-kPRED. **(a)** Scatterplot of score versus accuracy for the 15,544 query proteins of PDB30 dataset. (b) Scatterplot of scores for predicted versus actual PB sequences for the 15,544 query proteins of PDB30 dataset. Datapoints are coloured based on level of accuracy of predictions.

### Case studies

Here were considered the predictions for 5 specific cases to look at the strengths and limitations of the PB-kPRED algorithm namely in presence or absence of homologues of known structure. Results are reported in Table 6 and further described below. These case studies correspond to counter intuitive prediction instances where (i) prediction accuracy is high despite not having any close homologues and (ii) prediction accuracy is low despite having sequences of PBs of closely-related proteins in PENTAdb.

**Table 6.**
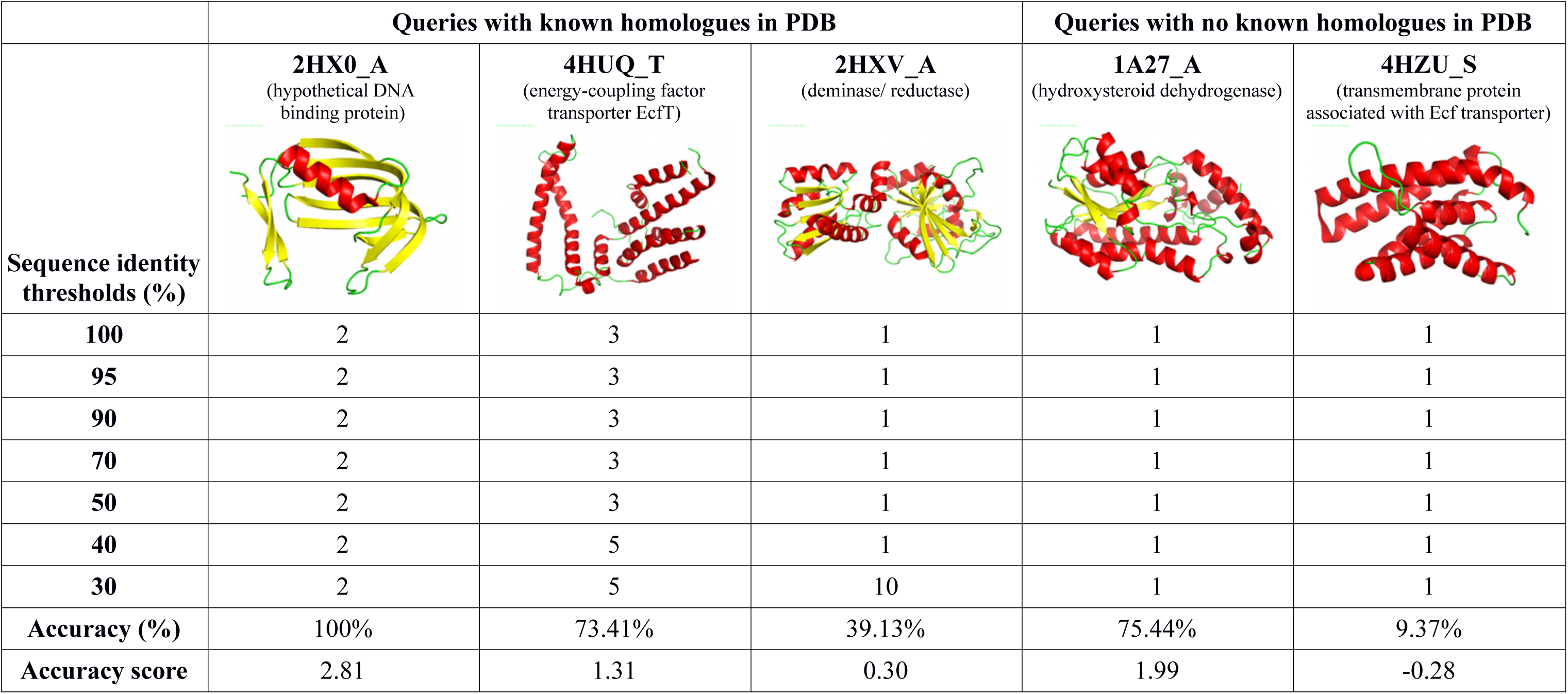
Impact of the availability of known homologues on the accuracy of PB-kPRED. Query PDB chains with known homologues and with no known homologues are featured. The *hybrid method with noise filtering* scheme for querying the database was applied for the prediction whereby the conditions were identical to experiment B1 as featured in Figure 3 and Table 6. The queries themselves were excluded from the database prior to prediction. Shown here are the accuracies of the predictions and the numbers of known homologues for different sequence identity thresholds.

|  | Queries with known homologues in PDB |  |  | Queries with no known homologues in PDB |  |
| --- | --- | --- | --- | --- | --- |
| Sequence identity thresholds (%) | **2HX0_A**<br>(hypothetical DNA binding protein) | **4HUQ_T**<br>(energy-coupling factor transporter EcfT) | **2HXV_A**<br>(deminase/ reductase) | **1A27_A**<br>(hydroxysteroid dehydrogenase) | **4HZU_S**<br>(transmembrane protein associated with Ecf transporter) |
| **100** | 2 | 3 | 1 | 1 | 1 |
| **95** | 2 | 3 | 1 | 1 | 1 |
| **90** | 2 | 3 | 1 | 1 | 1 |
| **70** | 2 | 3 | 1 | 1 | 1 |
| **50** | 2 | 3 | 1 | 1 | 1 |
| **40** | 2 | 5 | 1 | 1 | 1 |
| **30** | 2 | 5 | 10 | 1 | 1 |
| **Accuracy (%)** | 100% | 73.41% | 39.13% | 75.44% | 9.37% |
| **Accuracy score** | 2.81 | 1.31 | 0.30 | 1.99 | -0.28 |

Regarding prediction in employing information from homologues of known structure, three contrasting cases were studied. The first case relates to chain A of a hypothetical DNA binding protein from *Salmonella cholera* (PDB id 2HX0_A) which has a homologue from *Salmonella typhimurium* (PDB id 2NMU) that is 100% identical, 100% accuracy was achieved as shown by the high accuracy score of 2.81. Both structures aligned very well with a RMSD of 0.14 Å (Supplementary Figure 2a). The second case relates to an energy-coupling factor transporter transmembrane protein EcfT from *Lactobacillus brevis* (PDB id 4HUQ_T) which has two other “homologues” (PDB id 4RFS_T and 4HZU_T) that are 100% identical to the query. Here, the prediction accuracy is 73.4% only with an accuracy score of 1.31. 3-D structural alignment with these two “homologues” resulted in RMSDs 1.41 Å and 1.90 Å respectively displaying some structural variations (Supplementary Figure 2b) despite being 100% identical at the amino acid sequence level. These structural variations were due to rigid body movement. The third case is chain A of a pyrimidine deaminase / uracil reductase from *Thermotoga maritima* (PDB id 2HXV_A) which had only ten very distantly related proteins that shared less than 30% sequence identity in the PDB. Prediction rate is even lower here with accuracy reaching a value of 39.1% as shown by the low accuracy score of 0.30.

As for predictions in absence of homologues of known structure, two contrasting cases were studied. The first case is about a human hydroxysteroid dehydrogenase and the second case is a membrane protein associated with Ecf transporter from *Lactobacillus brevis*. The prediction performed quite well in the first case with an accuracy of 75.4% as shown by the high accuracy score of 1.99 while in the second case, the prediction almost completely failed with the accuracy of only 9.37% and also shown by the unfavorable accuracy score of −0.28.

### Implementation of the PB-kP*RED* methods as a web-tool

The PB-kPRED methodology has been implemented as a web-tool that is freely available to the community at http://www.bo-protscience.fr/kpred/. Both *majority rule* and *hybrid methods* without the *noise filtering scheme* for querying the database have been implemented. The tool provides a predicted PB sequence for each query amino acid sequence and also provides the *accuracy score* that serves as an *a posteriori* estimation of the prediction accuracy. In case the prediction score value is below −1, the prediction accuracy cannot be estimated and the user is notified. Users can provide multiple query protein sequences. All results are downloadable as FASTA formatted flat files. Optionally when submitting numerous query sequences, the user can provide an email address to which a notification will be sent when the job is completed.

## Discussion

The exponential growth in the structural knowledge of proteins has warranted the necessity of competent knowledge-based prediction algorithms for local structure prediction. At the level of short protein segments like pentapeptides, this increase in structural knowledge invariably brings with it an unprecedented signal to noise ratio for deciding on the most probable local conformations. Indeed, it is well established that similar pentapeptides can adopt different local conformations^3,33^. This is verified when the content of PENTAdb is inspected.

Hazout’s team along with defining the protein blocks also predicted the local structure in terms of PBs using a Bayesian approach^12^. They achieved an accuracy of 34.4% using a 15-residue window and this increased to 40.7% upon supplementing the Bayesian predictor with sequence profiles in the form of *sequence families*. In 2005, Etchebest and colleagues^26^ used a combination of statistical optimization procedure and improved sequence family data to bump up the accuracy to 48.7%. Incorporating secondary structure predictions from PSI-PRED into the PB prediction process did not contribute much to improve the accuracy *i.e* only 1% gain resulting in 49.9%. Machine learning techniques have also been used to predict protein local structure in terms of PBs. Support vector machine based methods like LOCUSTRA^20^, svmPRAT^22^ and SVM-PB-Pred^21^ achieve mean accuracies of 61.0%, 67.0% and 53.0% respectively. A dual layer neural network based prediction method achieved 58.5% accuracy^24^. The most refined version of the PB-kPRED method proposed here, i.e *hybrid method* with *noise filtering scheme*, outperformed most of the previously developed methods for PB prediction except for svmPRAT where it performed equivalently. Although all the methods evaluated their accuracies on non-redundant sets of proteins, an even comparison is hindered by difference in datasets, varying training regimes for the machine learning methods and different levels of sequence identity used as input in the prediction process. This motivated us to perform a battery of tests on the algorithm to estimate the prediction accuracy when incremental levels of sequence identities are made available in PENTAdb for the prediction (see Table 4). Importantly, to our knowledge, this is the first report of a querying scheme that dynamically filters out, on a per query basis, homologues at different cutoff values so that the portion of the PENTAdb that is made accessible for prediction is calculated on the fly. For the each of the 15,544 query sequences of PDB30, 16 experiments were performed amounting a total of 248,704 datasets building. This is computationally intensive and was performed using extensive MySQL querying. Thanks to the *noise filtering* strategy, PB-kPRED was able to efficiently weed out the noise present in the database due to redundancy and hence to narrow down the search in the database to find the most appropriate local structure for a given pentapeptide. Hence, filtering out from the database the pentapeptides from proteins that shared less than 30% sequence identity with the query indeed improves the prediction efficiency.

When the *majority method* and the *hybrid method* (Figure 5) were compared, two distinct clusters were noticed. Upon further investigating the reason for this distinct clustering, we note that, irrespective of the sequence identity cut-off, the points below the diagonal were found in more populated clusters while the points above the diagonal were found in least populated clusters Hence the *hybrid method* using the *noise filtering scheme* will perform better when there are some closely-related protein structures to look-up to in PENTAdb. In a real-life scenario, this will not be always the case. Indeed, proteins for which we want to predict the structure and which do not have any homologues even at 30% sequence identity are not so uncommon. This brings us to the conclusion that even though overall the *hybrid method* performs better, we cannot ignore the *majority rule method* all together.

Nonetheless, this method still has room for improvement as it can be seen from the values in Table 2. The list of all possible PBs reported by the PB-kPRED algorithm after querying PENTAdb database indeed shows that the good PB was present in more than 70% of the cases. However, owing to the scoring functions S1 and S2 (Figure 2), the decision rules implemented in both *majority rule* and *hybrid methods* failed to pick up these good PBs as predictions in several instances.

Interestingly, once the local backbone of a protein was predicted in the form of a PB sequence, we were able to provide an *a posteriori* assessment of how accurate was the prediction. The method used here to achieve this relied on the simple idea that successions of PBs should follow the rule that not all combinations of PBs would be allowed. This intuition turned out to be correct since there was a remarkable correlation between the score and the accuracy of the predictions. Noteworthy, the *accuracy scores* for actual (native) PB sequences are overwhelmingly distributed between +1 and +3, while poorly predicted PB sequences have scores below +1. This scoring of PB sequences could also serve as an indicator towards improving predictions. Because the calculation of the score of a PB sequence is very fast, one could imagine implementing a score-guided optimization procedure to climb the prediction accuracy gradient using Monte-Carlo or genetic algorithms for example.

The case studies documented in this work (see Table 6) indicate that the relationship between local structure predictability and the number of homologues of the query available in PDB are not very straightforward. Optimistically, in spite of not having any homologues, the PB-kPRED algorithm can perform a good prediction if the pentapeptides constituting the query adopt consensus local structures for the respective pentapeptides. Two such examples were provided but with contrasting outcomes, one achieving good accuracies and the other failing to predict correctly the PB sequence. Interestingly, the *accuracy scores* provided by our scoring function helped to reliably differentiate one prediction from the other. On the other hand, even if a query as multiple homologues in the PDB, its prediction accuracy will take a hit if the homologues are contrasting structural analogues of the query. For example, the activation of human pancreatic lipase involves considerable conformational transition in the form of a ‘lid movement’. The hypothetical prediction case when the query is the ‘lid open form’ and PENTAdb has pentapeptides from the ‘lid closed form’ would confuse the prediction algorithm despite both the forms of lipase being identical in amino acid sequences. Hence these case studies establish two take home messages: (i) there are exceptions to the general observation that the presence of homologues improves the prediction accuracy of PB-kPRED and (ii) the *accuracy score* used to evaluate the predictions is a reliable gauge for estimating the accuracy of the method as illustrated in Figure 6.

PB-kPRED web-server could form a vital link in the pipeline of PB based structure analysis tools. Namely, it can be bridged with PB-based fast structure comparison tools like iPBA^17^ and PBalign^16^ and help to mine for similar structures and map the fold space. It can also be used to predict the occurrence of structural motifs in protein sequences. Indeed, the alpha version of the server which was made available on-line earlier, has already been used by some research groups for the structural characterization of RNA binding sites in protein structures and predicting proteins sequences that contain RNA binding sites^34,35^ and also in predicting β-turns and their types^36^.

The web-based tool currently does not feature the *hybrid method* with *noise filtering scheme* because it would require running an instance of BLASTClust on every query. We plan to implement this functionality in a future improvement to the tool.

## Supplementary material

Additional methods file: additional_methods.pdf

Supplementary tables file: supplementary_tables.pdf

Supplementary figures file: supplementary_figures.pdf

## Acknowledgements

Authors thank Tristan Rialland and Sara Bachiri for providing technical support in the development of the PB-kPRED web server. This work was supported by the Région Réunion and the Fond Social Européen [grant no. 20131528] to IV. This work was in part supported by Conseil Régional des Pays de la Loire in the framework of GRIOTE project. AdB and FC acknowledge grants from the Ministry of Research (France), National Institute for Blood Transfusion (INTS, France), National Institute for Health and Medical Research (INSERM, France) and labex GR-Ex. The labex GR-Ex, reference ANR-11-LABX-0051 is funded by the program “Investissements d’avenir” of the French National Research Agency, reference ANR-11-IDEX-0005-02. AdB acknowledge supports by University Paris Diderot, Sorbonne, Paris Cité (France), FC acknowledge supports by Université de La Réunion, Faculty of Sciences and Technology. NS and AdB acknowledge to Indo-French Centre for the Promotion of Advanced Research / CEFIPRA for collaborative grant (number 5302-2). Research in NS laboratory is also supported by Department of Biotechnology, Government of India. NS is a J.C. Bose National Fellow.

